# *Caenorhabditis elegans* heterochromatin factor SET-32 plays an essential role in transgenerational establishment of nuclear RNAi-mediated epigenetic silencing

**DOI:** 10.1101/255562

**Authors:** Natallia Kalinava, Julie Zhouli Ni, Zoran Gajic, Helen Ushakov, Sam Guoping Gu

## Abstract

Epigenetic inheritance contributes fundamentally to transgenerational physiology and fitness. Mechanistic understanding of RNA-mediated chromatin modification and transgenerational epigenetic inheritance, which in *C. elegans* can be triggered by exogenous double-stranded RNA (exo-dsRNA) or facilitated by endogenous small interfering RNAs (endo-siRNAs), has mainly been limited to the post-initiation phases of silencing. Indeed, the dynamic process by which nuclear RNAi engages a transcriptionally active target, before the repressive state is stably established, remains largely a mystery. Here we found that the onset of exo-dsRNA-induced nuclear RNAi is a transgenerational process, and that establishment requires SET-32, one of the three putative histone methyltransferases (HMTs) that are required for H3K9me3 deposition at the nuclear RNAi targets. We also performed multigenerational whole-genome analyses to examine the establishment of silencing at endogenous targets of germline nuclear RNAi. The nuclear Argonaute protein HRDE-1 is essential for the maintenance of nuclear RNAi. Repairing a loss-of-function mutation in *hrde-1* by CRISPR restored the silencing of endogenous targets in animals carrying wild type *set-32*. However, for numerous endogenous targets, repairing the *hrde-1* mutation in a *set-32;hrde-1* double mutant failed to restore their silencing states in up to 20 generations after the *hrde-1* repair, using a similar genome editing approach. We found that despite a prominent role in the establishment of silencing, however, *set-32* is completely dispensable for the maintenance of silencing once HRDE-1-dependent gene repression is established. Our study indicates that: 1) establishment and maintenance of siRNA-guided transcriptional repression are two distinct processes with different genetic requirements; and 2) the rate-limiting step of the establishment phase is a transgenerational, chromatin-based process. In addition, our study reveals a novel paradigm in which a heterochromatin factor primarily functions to promote the establishment of transgenerational silencing, expanding mechanistic understanding of the well-recognized role of heterochromatin in epigenetic maintenance.

## Introduction

The term RNA interference (RNAi) originally refers to the phenomenon of exo-dsRNA-triggered gene silencing (Fire et al., 1998; Kennerdell and Carthew, 1998), in which target messenger RNA (mRNA) is degraded by an siRNA-associated AGO protein, resulting in post-transcriptional gene silencing (PTGS) (Elbashir et al., 2001; Hammond et al., 2000; Hammond et al., 2001; Tuschl et al., 1999). In addition to this mechanism, siRNAs can also target a gene for transcriptional gene silencing (TGS) in plants, fungi, and animals (Martienssen and Moazed, 2015; Pezic et al., 2014; Sienski et al., 2012; Wassenegger, 2000). We will use the terms classical RNAi and nuclear RNAi to refer to PTGS and TGS, respectively. Classical RNAi results in rapid degradation of exogenous dsRNA and homologous single-stranded transcripts, while long term, stable silencing of transposons and other types of repetitive genomic elements is facilitated by nuclear RNAi.

In *C. elegans*, nuclear RNAi effects include histone modifications (H3K9me3 and H3K27me3) and transcriptional repression at endo-siRNA-targeted loci (Buckley et al., 2012; Gu et al., 2012; Guang et al., 2010; Mao et al., 2015). The study of the endogenous silencing events, which provide a rich source of targets to study the physiological functions and underlying mechanisms, is further complemented by highly manipulatable approaches using exogenous triggers such as dsRNA or transgenes (Ashe et al., 2012; Gu et al., 2012; Guang et al., 2010; Leopold et al., 2015; Minkina and Hunter, 2017; Shirayama et al., 2012). Exo-dsRNA-induced nuclear RNAi relies on the upstream steps of classical RNAi for siRNA biogenesis (Grishok et al., 2000; Gu et al., 2012), but also requires nuclear RNAi-specific protein factors, including the germline-specific nuclear AGO protein WAGO-9/HRDE-1 (Akay et al., 2017; Ashe et al., 2012; Buckley et al., 2012; Guang et al., 2010; Shirayama et al., 2012; Spracklin et al., 2017; Weiser et al., 2017). The germline nuclear RNAi pathway is the core effector for various transgenerational silencing phenomena in *C. elegans* (Alcazar et al., 2008; Ashe et al., 2012; Bagijn et al., 2012; Burkhart et al., 2011; Gu et al., 2012; Leopold et al., 2015; Minkina and Hunter, 2017; Shirayama et al., 2012). In the case of exo-dsRNA-induced heritable RNAi, the heterochromatin response, as well as the silencing effect, can persist for multiple generations after the initial dsRNA exposure (administered by feeding or injection) has been ceased (Alcazar et al., 2008; Ashe et al., 2012; Buckley et al., 2012; Grishok et al., 2000; Gu et al., 2012; Mao et al., 2015), providing a highly tractable system to study the transgenerational epigenetic inheritance of silencing.

The germline nuclear RNAi pathway plays an important role in maintaining genome stability (Bagijn et al., 2012; McMurchy et al., 2017; Ni et al., 2014) and is essential for germline development when *C. elegans* is under heat stress (Ashe et al., 2012; Buckley et al., 2012; Ni et al., 2016; Weiser et al., 2017). Based on the mapping of HRDE-1-associated endo-siRNAs, a diverse set of genomic regions are putative targets of the germline nuclear RNAi pathway (Buckley et al., 2012). We previously further refined the putative targets by identifying ones that lose H3K9me3, transcriptional repression, or both in mutant animals that lack HRDE-1 (Ni et al., 2014). We refer to these targets as the exemplary endogenous targets of germline nuclear RNAi, which include LTR retrotransposons, other types of repetitive DNA, as well as some protein-coding genes. Interestingly, regions with germline nuclear RNAi-dependent heterochromatin (GRH) and regions with germline nuclear RNAi-dependent transcription silencing (GRTS) only partially overlap (Ni et al., 2014).

H3K9me3 deposition at either exo-dsRNA target genes or the endogenous HRDE-1 targets requires multiple H3K9 methyltransferases, MET-2, SET-25, and SET-32 (Kalinava et al., 2017; Mao et al., 2015; Spracklin et al., 2017). Studies using different experimental setups and target genes have reported that H3K9 HMTs are required for the siRNA-guided epigenetic silencing effects in some cases (Ashe et al., 2012; Minkina and Hunter, 2017; Spracklin et al., 2017), but are dispensable in others (Kalinava et al., 2017; Lev et al., 2017; Minkina and Hunter, 2017). *met-2* mutant animals even exhibit the phenotype of enhanced heritable RNAi (Lev et al., 2017). We recently showed that H3K9me3 can be decoupled from transcriptional repression in nuclear RNAi (Kalinava et al., 2017). Abolishing the H3K9me3 deposition by mutating all the three H3K9 HMT genes does not lead to any derepression at the endogenous targets, nor does it cause any defects in transcriptional silencing or inheritance of silencing at an exo-dsRNA target gene. This argues against a simple model in which H3K9me3 plays a direct or dominant role in transcriptional repression at the nuclear RNAi targets. However, our previous study at endogenous targets only examined the requirement of H3K9 HMTs in maintenance of already established silencing states. It is unknown whether the H3K9 HMTs are required for de-novo silencing of the endogenous targets from a de-repressed state. For the assays of exo-dsRNA-induced nuclear RNAi, worms with several generations of exo-dsRNA exposure were used to achieve the steady-state level of repression (Kalinava et al., 2017). Therefore, the roles of H3K9 HMTs in the onset of nuclear RNAi were unknown. In this study, we investigated the transgenerational kinetics of nuclear RNAi-mediated silencing and H3K9 HMTs’ roles in this process.

## Results

### The onset of nuclear RNAi-mediated silencing in wild type animals occurs one generation after the exo-dsRNA exposure

To investigate the transgenerational onset of nuclear RNAi, we performed a five-generation RNAi experiment targeting *oma-1* (Fig. 1A), a non-essential germline gene (Lin, 2003) that has been routinely used to study nuclear RNAi and epigenetic inheritance of silencing. The *oma-1* dsRNA feeding began at the synchronized L1 larval stage of the first generation and continued for another four generations. Young adult worms were collected for analysis at each generation. Adult worms fed on *E. coli* OP50 (dsRNA-) were used as the control.

**Figure 1.**
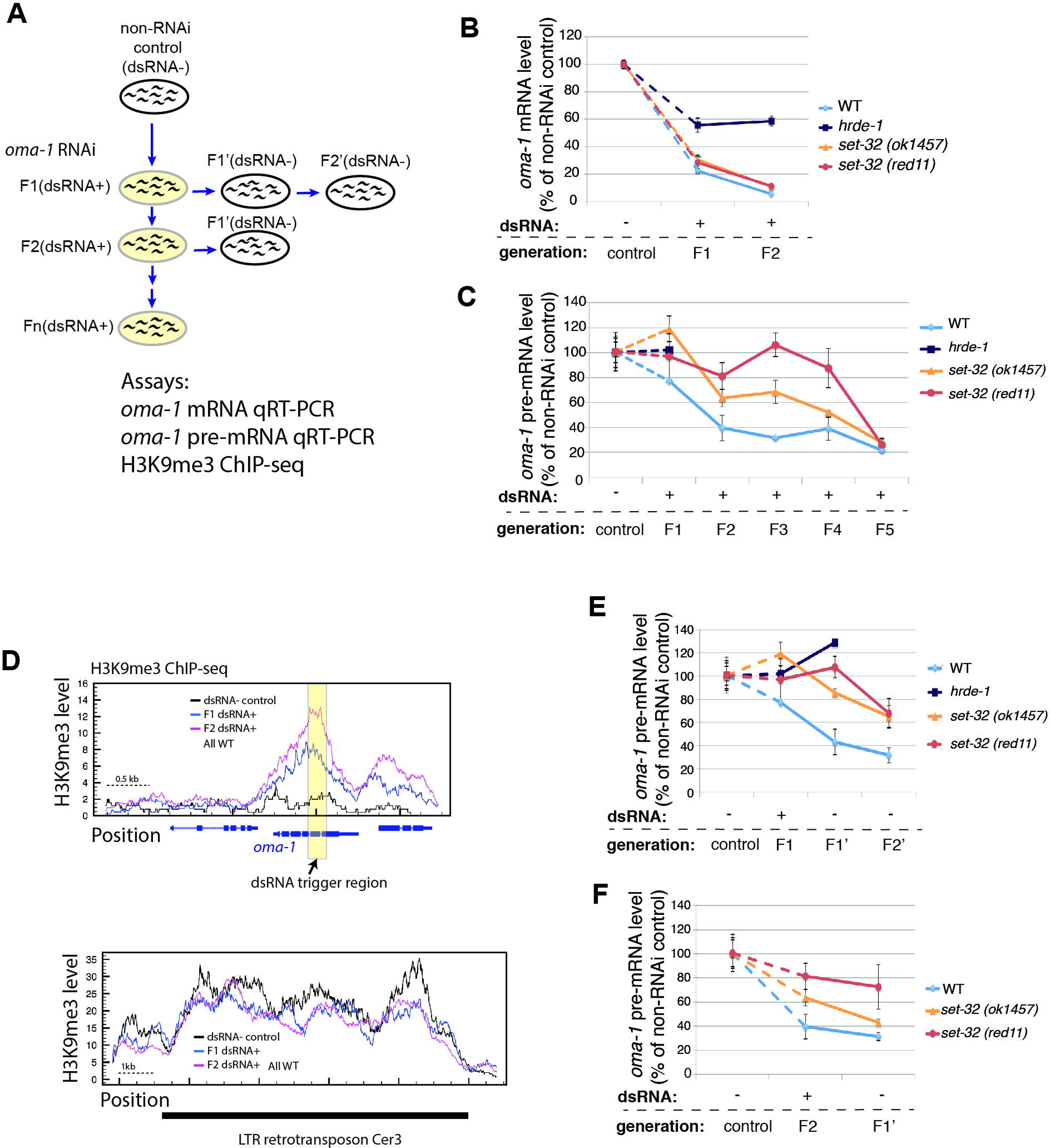
Multigenerational analysis of dsRNA-triggered RNAi at *oma-1.* (A) A schematic of the experiment showing that adult animals fed on *oma-1* dsRNA-expressing *E. coli* for one to five generations (F1-F5[dsRNA+]) and the inheritance generations fed on OP50 (F1’ and F2’[dsRNA−]) were collected for the analysis. (B) *oma-1* mRNA qRT-PCR analysis of the control (dsRNA−), F1(dsRNA+), and F2(dsRNA+) samples. (C) *oma-1* pre-mRNA qRT-PCR analysis of the control (dsRNA−) and F1-F5(dsRNA+) samples. (D) H3K9me3 ChIP-seq coverage plots in the *oma-1* locus (the top panel, RNAi target) and LTR retrotransposon Cer3 (the bottom panel), Cer3, used as a control locus here, is an endogenous HRDE-1 target with a high level of H3K9me3, which is not affect by *oma-1* RNAi. Wild type animals of control (dsRNA−), F1 (dsRNA+), and F2 (dsRNA+) were used. (E) *oma-1* pre-mRNA qRT-PCR analysis of the progeny (F1’[dsRNA−] and F2’[dsRNA−]) of the F1(dsRNA+) animals. (F) *oma-1* pre-mRNA qRT-PCR analysis of the progeny (F1’[dsRNA−]) of the F2(dsRNA+) animals. The values for the control, F1(dsRNA+), and F2(dsRNA+) samples from (C) are also used in (E) and (F) for comparison.

To examine the nuclear RNAi-mediated silencing at *oma-1*, we measured the *oma-1* pre-mRNA levels by using qRT-PCR, with random hexamer oligoes as the RT primer and an *oma-1* intron-specific and an exon-specific primer for the qPCR. To examine the combined effects of classical RNAi and nuclear RNAi at *oma-1*, we measured the *oma-1* mRNA levels using the oligo-dT as the RT primer and two exon-specific primers for the qPCR. We found that *oma-1* mRNA was largely depleted by RNAi at the second generation in wild type animals, down to 5% of the control mRNA level (Fig. 1B). The *oma-1* pre-mRNA at the second generation was reduced to a similar level in the later generations (Fig. 1C), indicating that the transcriptional repression reached the steady-state level at the second generation in the wild type animals.

During the first generation of the dsRNA exposure, the wild type animals showed more profound repression of *oma-1* at the mRNA level than at the pre-mRNA level. *oma-1* mRNA was reduced to 22% of the control level, indicating that the classical RNAi occurred robustly in the first generation (Fig. 1B). In contrast, *oma-1* pre-mRNA level was only reduced to 77% of the control level (Fig.1C) (no detectable loss of *oma-1* pre-mRNA in F1(dsRNA+) worms was observed in the second replica, Fig. S1), much higher than those observed in the subsequent generations. This result indicates that in wild type animals exo-dsRNA triggers a robust classical RNAi-mediated silencing in the first generation, while the onset of a robust nuclear RNAi occurs one generation after the initial dsRNA exposure.

We also examined H3K9me3 at the *oma-1* locus by performing H3K9me3 ChIP-seq using the F1(dsRNA+) and F2(dsRNA+) wild type animals, as well as the dsRNA- control. We found that both F1(dsRNA+) and F2(dsRNA+) animals exhibited high levels of H3K9me3 in *oma-1* gene (Fig. 1D), and the levels were similar to our previously reported level in animals treated with *oma-1* dsRNA for 3-4 generations (Kalinava et al., 2017). Although the F1(dsRNA+) sample had a lower H3K9me3 signal in *oma-1* than the F2(dsRNA+) sample, the H3K9me3 difference was smaller (F2/F1=1.4) than the difference in the transcription repression between these two samples (F2/F1=3.0).

We also examined the *oma-1* mRNA levels in *hrde-1* mutant animals that were exposed to the *oma-1* dsRNA for one and two generations. As expected, there was a lack of repression at the pre-mRNA level in the first generation (Fig. 1C). We observed approximately 40% reduction in *oma-1* mRNA in both the first and second generations compared to the control (dsRNA- worms) (Fig. 1B). This result suggests that the classical RNAi, which is active in the *hrde-1* mutant (Buckley et al., 2012; Kalinava et al., 2017), can compensate the transcriptional silencing defect caused by the *hrde-1* mutation. However, such compensation appeared to be partial possibly due to the robust transcription of *oma-1* in *hrde-1* mutant animals.

### set-32 mutation results in a multigenerational delay in the establishment of exo-dsRNA-induced transcriptional repression

To investigate whether any of the H3K9 HMTs plays a role in the initiation of transcriptional repression, we performed a multigenerational *oma-1* RNAi experiment using *set-32, met-2*, and *set-25* single mutant strains, as well as the *met-2 set-25* double mutant. Two different *set-32* mutant alleles were used in this study. One is the *set-32(ok1457)* allele which results in a 156 amino acids in-frame deletion (position 50-205, before the SET domain) (Consortium, 2012) and has been used in previous studies (Ashe et al., 2012; Kalinava et al., 2017). The *set-32(ok1457)* allele does not eliminate the SET domain which contains the catalytic center for the HMT reaction. To determine whether *set-32(ok1457)* is inactive in depositing H3K9me3 at the target chromatin, we generated a second allele, *set-32(red11)*, by CRISPR that lacks the SET domain. We verified that both alleles had very similar defects in dsRNA-induced H3K9me3 at *oma-1*. As previously observed for the *set-32(ok1475)* mutant (Kalinava et al., 2017), *set-32(red11)* mutant animals exhibited no defect in heritable RNAi after approximately four generations of exo-dsRNA exposure (Fig. S2A and S2B). *oma-1* mRNA was robustly silenced by RNAi in animals carrying either of the two *set-32* mutant alleles at the first generation of the dsRNA exposure, similar to wild type animals (Fig. 1B). The onset of nuclear RNAi, measured by pre-mRNA, was much slower in *set-32* mutant than the wild type animals. Both *set-32* mutant alleles generally had higher *oma-1* pre-mRNA than wild type animals in the first four generations (Fig. 1C and Fig. S1). It was at the fifth generation when the repression reached the steady-state level observed in the wild type animals (Fig. 1C and Fig. S1). Therefore, *set-32* mutation causes a multigenerational delay in dsRNA-triggered transcriptional repression. We also tested *met-2, set-25,* and *met-2 set-25* mutant animals. All three mutants exhibited wild type-like pre-mRNA repression at the first and second generations of the dsRNA exposure (Fig. S1B). These results suggest that, although both SET-32 and MET-2 SET-25 contribute to the H3K9me3 at the RNAi target, their functions are not equivalent. SET-32-dependent activity, but not MET-2 SET-25-dependent activity, promotes the onset of exo-dsRNA-triggered transcriptional repression.

To examine silencing inheritance induced by a shorter duration of dsRNA exposure (one or two generations as opposed to five), we cultured the progeny of dsRNA-exposed animals on OP50 plates that lacked *oma-1* dsRNA for one or two generations (referred to as inheritance generations) (Fig. 1A). For wild type animals, both one-generation and two-generation dsRNA exposures resulted in robust repression at the pre-mRNA level in the inheritance generations, with a transgenerational kinetics that is similar to continuous dsRNA exposure (Fig. 1E and F). Therefore, the one-generation delay in nuclear RNAi in wild type animals is not due to an insufficient duration of dsRNA exposure. For *set-32* mutants, we again observed delayed nuclear RNAi in the inheritance generations (Fig. 1E and F), confirming the role of SET-32 in promoting the onset of nuclear RNAi-mediated silencing.

### SET-32 is required for establishment of silencing for some of the endogenous targets of the germline nuclear RNAi pathway

To examine the establishment of silencing at the endogenous targets, we used CRISPR-mediated gene conversion to repair the loss-of-function *hrde-1(tm1200)* mutation in worms that had carried the *hrde-1* mutation for more than 20 generations. We wished to know how fast the repressive states can be established at the endogenous nuclear RNAi targets when the HRDE-1 activity is restored. Besides the CRISPR method, a wild type *hrde-1* allele can also be introduced to the *hrde-1* mutant strain via a genetic cross. We did not choose this method because genetic crosses will introduce epigenetically repressed alleles of the endogenous targets, as well as a foreign siRNA population.

To repair the *hrde-1* mutation using CRISPR, a mixture of *hrde-1*-targeting Cas9 ribonuclease complex and repair template DNA was injected into the gonads of hermaphrodite adults (Fig. 2A). The wild type *hrde-1* allele generated by gene conversion was annotated as *hrde-1*(+^R^) to distinguish it from the naïve wild type *hrde-1* allele *(hrde-1[+], i.e.,* never experienced any mutation in the gene). Self-fertilized progeny with the *hrde-1*(+^R^/−) (F1) and *hrde-1*(+^R^/+^R^) (F2) genotypes were identified by PCR and Sanger sequencing. We then collected later generations of the homozygous *hrde-1*(+^R^) adults and examined the abundance of the transcripts of the endogenous targets. The F3 generation was the earliest one that we were able to collect for the assay because the F1 and F2 progeny were used for genotyping. We performed qRT-PCR analysis for two endogenous targets, of which, both established their repressive states in the F3 and later generations in a wild type background (Fig. 2B and C).

**Figure 2.**
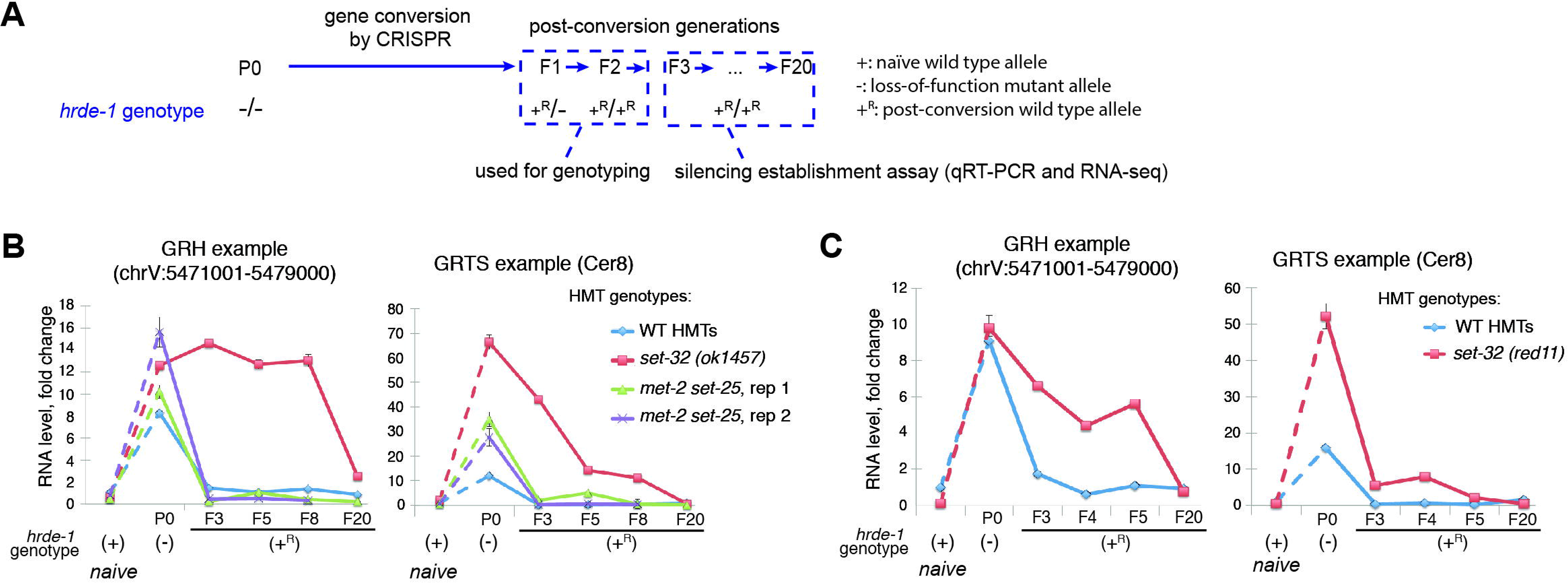
Silencing establishment assay of endogenous HRDE-1 targets. (A) A schematic of the silencing establishment assay using CRISPR-Cas9-mediated gene conversion of the *hrde-1(tm1200)* mutation to the wild type sequence. (B and C) qRT-PCR analysis for two endogenous HRDE-1 targets at pre‐ and post-conversion generations, as well as samples with the identical genotypes but without experiencing the *hrde-1* mutation (naïve samples). For *met-2 set-25*, results from two independent gene-conversion events were shown in (B). The GRH locus used in this analysis is chrV:5471001-5479000. The GRTS locus used in this analysis is LTR retrotransposon Cer8, located at chrV:5179680-5191222.

To test if H3K9 HMTs are required for establishment of silencing at the endogenous targets, we generated *set-32;hrde-1* and *met-2 set-25 hrde-1* mutant strains. In one set of experiments, we collected F3, F5, F8, and F20 populations after the gene conversion. In another set, we collected F3, F4, F5, and F20 populations. In the first set of the experiments, two independent lines of *met-2 set-25 hrde-1*[+^R^/+^R^] worms were followed and both fully established the repressive states by the F3 generation and in all subsequent generations for the two tested targets (Fig. 2C). In contrast, both *set-32* alleles exhibited a delay in the establishment of silencing for at least eight generations. Full repressions of these targets were reached at the F20 generation (Fig. 2B and C). We confirmed that the early generations of the *hrde-1*(+^R^) animals had similar *hrde-1* mRNA expressions as the *hrde-1*(+) animals (Fig. S3). Therefore, the delayed establishment of silencing is not due to a lower hrde-1 mRNA expression in the early post-conversion generations.

To examine the establishment of silencing at the whole-genome level, we performed RNA-seq for the post-conversion populations (*set-32;hrde-1*[+^R^/+^R^] and *met-2 set-25 hrde-1*[+^R^/+^R^]), as well as animals of the identical genotypes but with the naïve wild type hrde-1 allele (*set-32;hrde-1*[+/+] and *met-2 set-25 hrde-1*[+/+]). We sequenced rRNA-depleted total RNA of each sample in order to include both polyadenylated RNA and non-polyadenylated RNA. For the post-conversion samples, we performed RNA-seq for the F4, F5, and F20 populations for *hrde-1*(+^R^), F3, F5, and F20 for *met-2 set-25 hrde-1*(+^R^), F5, F8, and F20 for *set-32(ok1457) hrde-1*(+^R^), and F4, F5, and F20 for *set-32(red11) hrde-1*(+^R^). Our whole genome analysis indicated that repairing *hrde-1* mutation in animals carrying the wild type H3K9 HMTS resulted in establishment of silencing for all of the endogenous targets as early as in the F4 post-conversion population (Fig. 3A and E). Remarkably, a significant fraction of the endogenous targets remained actively expressed even in *set-32 hrde-1*(+^R^) F20 samples for both *set-32(red11)* and *set-32(ok1457)* alleles (Fig. 3C-E). These targets are referred to as putative irreversible targets. After examining the RNA-seq profiles individually, we identified four exemplary irreversible targets, which include two LTR retrotransposons (Cer9 and Cer12), a putative protein-coding gene c38d9.2 in chromosome V, and an approximately 5 kb region in chromosome II (containing two uncharacterized putative protein-coding genes f15d4.5 and f15d4.6) (Fig. 3F and Fig. S4B-E). All of these loci became transcriptional activated in *hrde-1* and *set-32;hrde-1* mutants, but remained silent in *set-32* mutant (with naïve wild type *hrde-1*), despite the losses of H3K9me3 at these regions (Fig. 3F and Fig. S4B-E). These data confirm our previously finding that SET-32 is dispensable for the maintenance of silencing at the native HRDE-1 targets (Kalinava et al., 2017). These irreversible targets represent a bona fide epigenetic phenomenon in which two genetically identical populations (cultured in the same conditions and harvested at the same developmental stage) exhibit different gene expression profiles.

**Figure 3.**
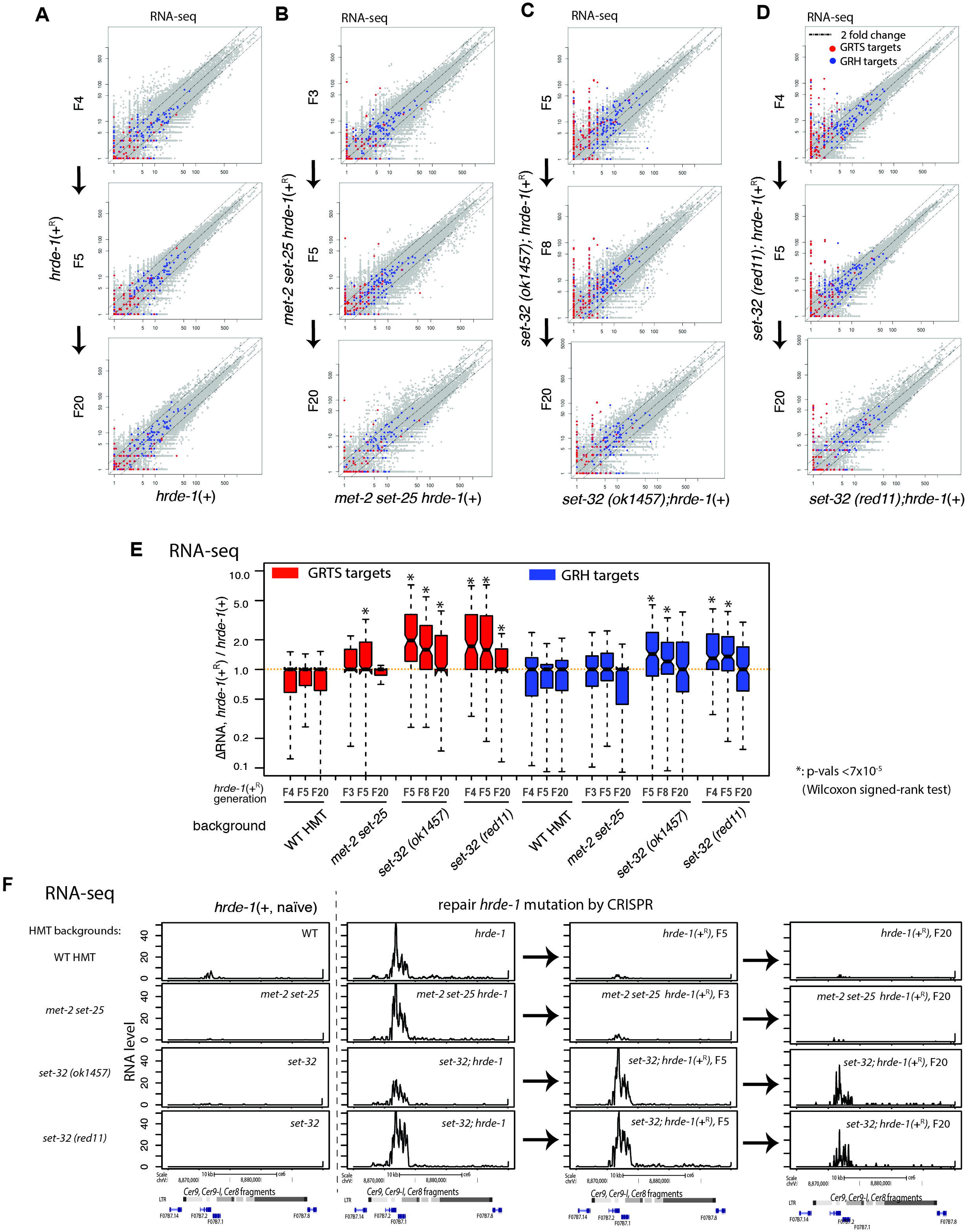
RNA-seq analysis of silencing establishment at endogenous HRDE-1 targets. (A-D) Scatter plot analysis of normalized RNA-seq counts for all 1kb genomic regions comparing animals in the post-conversion generations with animals of the same genotype but without experiencing *hrde-1* mutation (naïve samples). All samples in (A) carried wild type H3K9 HMT genes, (B) carried *met-2 set-25* double mutations, (C) carried *set-32(ok1457)*, and D carried *set-32(red11)* single mutation. Exemplary endogenous HRDE-1 targets were highlighted with germline nuclear RNAi-dependent transcriptional silencing (GRTS) and heterochromatin (GRH) regions colored in red and blue, respectively. (E) Boxplot analysis of the changes in RNA-expression of GRTS or GRH regions between post-conversion animals with corresponding naïve samples. Statistically significant changes were indicated by asterisks. (F) An exemplary endogenous HRDE-1 target (LTR retrotransposon Cer9) that is defective in silencing establishment in the *set-32* mutant for at least 20 generations, but not in the *met-2 set-25* mutant. Normalized total RNA-seq signals were plotted. Columns from left to right are the naïve samples *(hrde-1[+])*, pre-conversion samples *(hrde-1[−])*, and F5 and F20 post-conversion samples *(hrde-1[+^R^])*. Different genetic backgrounds (WT or various H3K9 HMT mutants) are in different rows.

We found that *met-2 set-25* are required for the establishment of silencing in only one of the four irreversible targets, *c38d9.2* (Fig. S4D). Our global analysis indicated that most of the endogenous targets established the silencing states as early as in the F3 *met-2 set-25 hrde-1*(+^R^) populations (Fig. 3B and 3E).

We note that majority of the irreversible targets in *set-32* mutant belong to the GRTS (germline nuclear RNAi-dependent transcriptional silencing) regions. In contrast, most of the GRH (germline nuclear RNAi-dependent heterochromatin) regions restored the transcriptional repression by the F3 post-repair generation in the *set-32* mutant background (Fig. 3C-E). We also performed the silencing establishment assay using the *set-32;met-2 set-25 hrde-1* mutant strain, and obtained similar results as the *set-32;hrde-1* mutant (Fig. S4 and data not shown). Taken together, our results indicate that SET-32 is required for the establishment of the repression states of some endogenous HRDE-1 targets. Although *met-2 set-25* mutant animals had a much weaker phenotype than *set-32* in this assay, we cannot rule out that MET-2 and SET-25 are required for the establishment of silencing in the first and second post-conversion generations, which were precluded in the assay due to the screening procedure after CRISPR.

To explore the role of endo-siRNAs in the establishment of silencing, we performed sRNA-seq analysis for the pre-conversion samples, samples of the F5 and F20 post-conversion generations, and the matching samples with the naïve wild type *hrde-1* allele. We found that the post-conversion *set-32 hrde-1*(+^R^) populations had abundant endo-siRNA profiles at the irreversible targets (Fig. S4B-E), arguing against the possibility that the delayed establishment phenotype is caused by a lag in endo-siRNA expression. This data also suggests that SET-32 functions in a step upstream to the siRNA-dependent activities.

### H3K9 HMTs are required to maintain the repressive states for some endogenous HRDE-1 targets in *hrde-1* mutant animals

We performed Pol II ChIP-seq and RNA-seq analysis for various compound mutants including *set-32;hrde-1, met-2 set-25 hrde-1,* and *set-32;met-2 set-25 hrde-1*. As expected, the endogenous HRDE-1 targets were derepressed in these compound mutants (Fig. S5A and B). Intriguingly, for a small subset of the targets, the *hmt; hrde-1* compound mutants exhibited much higher RNA expression and Pol II occupancy than the *hrde-1* single mutant (Fig. 4A and S5), indicating that the loss of H3K9me3 further enhances the *hrde-1* mutation’s defects in transcriptional repression for these targets. Such enhancement is unexpected because *set-32, met-2 set-25, set-32;met-2 set-25* mutant animals have no silencing defects at the endogenous targets, as reported previously (Kalinava et al., 2017) and shown again using the new data generated in this study (Fig. S5). An exemplary endogenous target with such enhanced derepression is the LTR retrotransposon Cer3 (Fig 4C-F). Either *set-32* single or *met-2 set-25* double mutations, when combined with the *hrde-1* mutation, drastically increased the Pol II occupancy at Cer3 and Cer3 transcripts. Therefore, the enhanced derepression occurs at the transcriptional level. The enhanced derepression caused by different H3K9 HMTs appears to be additive because the quadruple mutant of *set-32;met-2 set-25 hrde-1* showed the highest Cer3 expression (Fig. 4D-E). Interestingly, Cer3 is not an irreversible target (Fig. 4F). These results indicate that, for a small subset of the endogenous targets, H3K9me3 HMTs are required for the maintenance of repression. However, such requirements are only apparent when HRDE-1 activity is compromised. We note that, for the majority of the endogenous HRDE-1 targets, the combined mutations in *H3K9* HMT genes and *hrde-1* had no additive effect of derepression compared to the *hrde-1* single mutant (Fig. 4B and S5).

**Figure 4.**
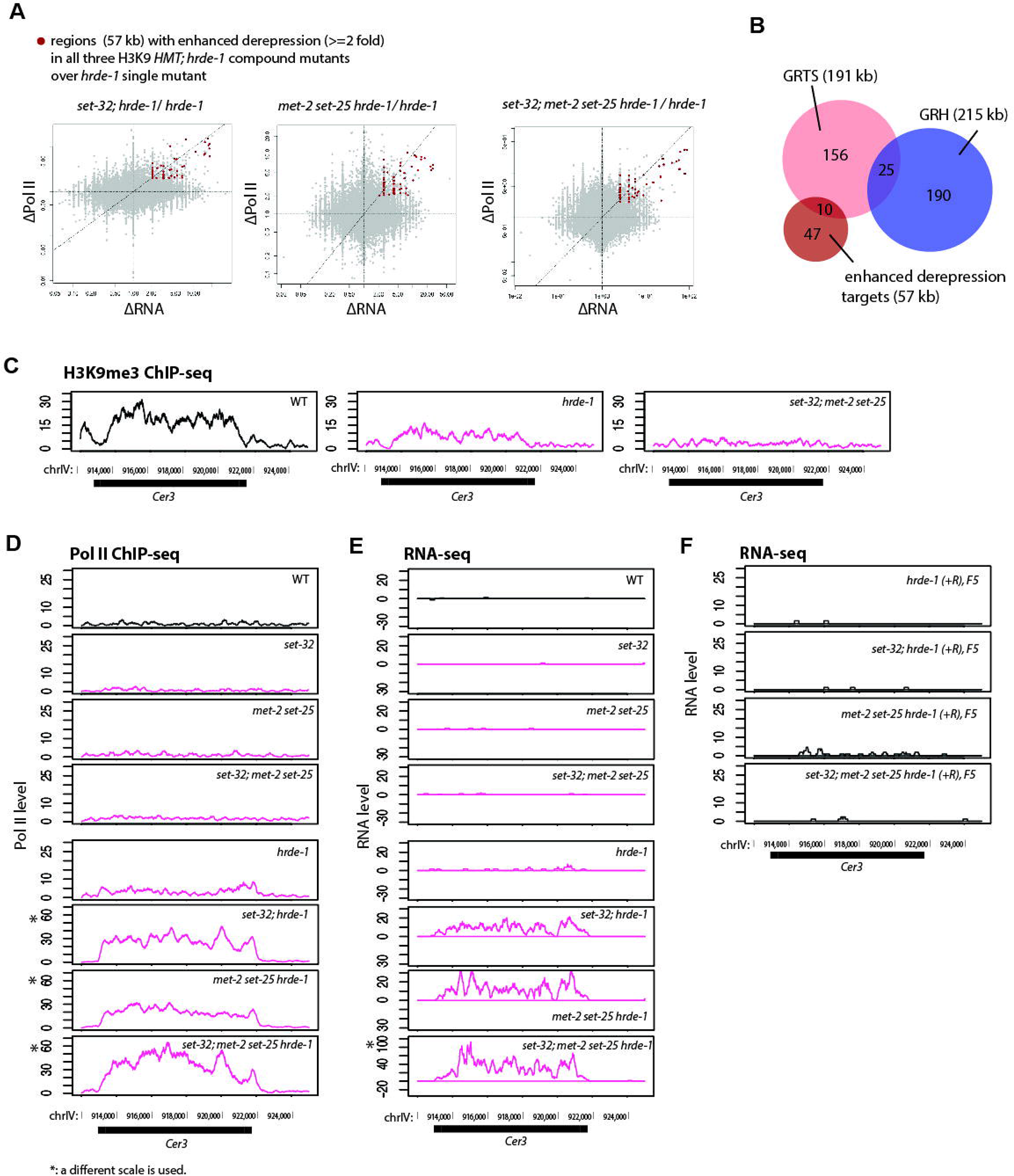
Enhanced silencing defects in the endogenous HRDE-1 targets caused by the combination of H3K9 HMT and *hrde-1* mutations. (A) Scatter plots with changes in RNA-seq signals between a compound mutant and *hrde-1* single mutant plotted in the X-axis and changes in Pol II ChIP-seq signals plotted in the Y-axis for all 1 kb genomic regions. Regions with ≥2-fold increases in both RNA expression and Pol II occupancy in all three *H3K9 hmt;hrde-1* compound mutants over *hrde-1* single mutant were marked with red dots. (B) Venn diagram showing the relationship between GRTS, GRH, and regions with enhanced derepression in the three *H3K9 hmt;hrde-1* compound mutants. (C-E) The H3K9me3 ChIP-seq (data from (Kalinava et al., 2017)), Pol II ChIP-seq (S2 phosphorylated), and RNA-seq coverage plots at the LTR retrotransposon Cer3 locus in wild type and various mutant animals. Panels marked with an asterisk (*) use a different scale to accommodate the enhanced expression levels observed for the genotype. (F) RNA-seq profiles at Cer3 for post-conversion samples with wild type H3K9 HMT genes or various H3K9 HMT mutations.

## Discussion

### The establishment and maintenance phases of germline nuclear RNAi in *C. elegans*

H3K9me3 is an evolutionarily conserved nuclear RNAi effect in fungi, plants, and animals. In *C. elegans* this effect is highly specific and prominent at target loci. Furthermore, it is transgenerationally heritable and linked to the germline immortality (Ashe et al., 2012; Buckley et al., 2012; Burton et al., 2011; Gu et al., 2012; Shirayama et al., 2012; Weiser et al., 2017). H3K9me3 at nuclear RNAi targets requires three HMTs: MET-2, SET-25, and SET-32 (Kalinava et al., 2017; Mao et al., 2015; Spracklin et al., 2017). Strikingly, we previously found that a complete loss of H3K9me3 in the mutant worms that lack the three HMTs did not cause any defect in transcriptional repression at the endogenous targets (Kalinava et al., 2017). The experimental setup in the previous study was designed to examine the maintenance of silencing and precluded us from studying the establishment phase. For a newly inserted transposon that is transcriptionally active, the establishment phase proceeds the maintenance phase and two phases may involve different mechanisms.

Exo-dsRNA-triggered nuclear RNAi provides an inducible system in which to study the onset, maintenance, and inheritance of epigenetic silencing at genes that are normally actively expressed. The previous discovery of a transgenerational delay in the onset of H3K9me3 at the target gene (Burton et al., 2011; Gu et al., 2012) suggests that the establishment of nuclear RNAi is a gradual process. The kinetics of transcriptional repression during the early generations of dsRNA exposure have not been examined before this study.

In this study, we examined the establishment phase for both endo-siRNA-guided and exo-dsRNA-triggered nuclear RNAi. Based on the results of this and our previous work (Kalinava et al., 2017), we propose a model in which the establishment and maintenance phases of nuclear RNAi have the following distinctions (Fig. 5).

**Figure 5.**
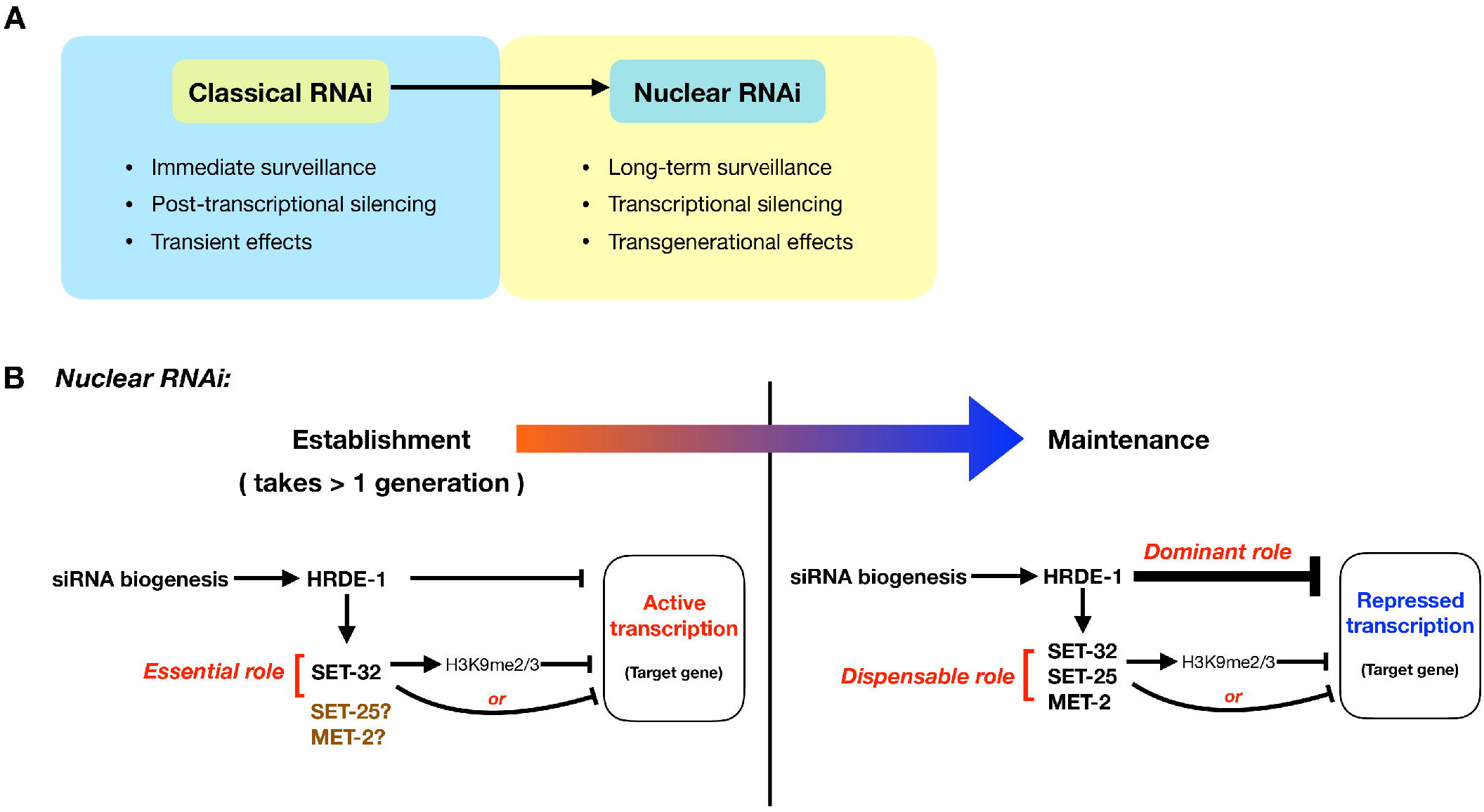
A schematic model that highlights the functional and mechanistic differences between (A) classical RNAi and nuclear RNAi, and (B) the establishment and maintenance phases of nuclear RNAi.

(1) The transcriptional state of the target locus. At the maintenance phase, the target is stably repressed and kept in a heterochromatic state. In contrast, the establishment phase begins with an actively transcribed and euchromatic state at the target locus. Converting from an active state to a stably repressed state requires removal of euchromatic marks and the deposition of heterochromatic marks. Therefore, the duration of the onset of silencing may depends on the relative strengths of the two opposing forces.

(2) Silencing mechanism. We propose that nuclear RNAi is a composition of multiple distinct silencing mechanisms. One of these mechanisms is a H3K9me3-independent silencing mechanism, which is likely the main source of silencing during the maintenance phase. The active transcription during the establishment phase may pose a strong force against the H3K9me3-independent silencing mechanism. Such antagonistic interactions between euchromatin (H3K36me3) and heterochromatin have been shown to occur in the *C. elegans* germline (Gaydos et al., 2012; Weiser et al., 2017). It is conceivable that SET-32-depedent H3K9me3 promotes the onset of silencing, either by providing an additional force of repression (H3K9me3-dependent in this case) or by enhancing the efficacy of H3K9me3-independent silencing mechanism. However, given enough time, H3K9me3-independent mechanisms can eventually establish silencing in the absence of SET-32. Although the prolonged lag in silencing is tolerated by the mutant worms at the organismal level in the laboratory condition, this molecular defect may create a window for the transposition of a newly invaded mobile DNA, and therefore it is likely to be selected against in a wild population.

The establishment phase of an epigenetic silencing phenomenon is often difficult to study because of its transient nature. The *set-32* mutation slows down the onset of silencing, and therefore provides an expanded window to dissect the extraordinarily complex process.

Although MET-2, SET-25, and SET-32 all contribute to the H3K9me3 at the nuclear RNAi targets, SET-32 and MET-2 SET-25 are not equivalent in their role in silencing establishment. This may due to differential expression of these H3K9 HMTs. Alternatively, the functional difference could be due to certain unknown H3K9me3-independent activities associated with individual enzymes. Future study is required to investigate these possibilities.

**What is the functional significance of the one-generation delay in exo-dsRNA-induced nuclear RNAi?** Although exo-dsRNA was used as an artificial means to induce RNAi in this study, it is akin to a situation in which host animals encounter viral nucleic acids. We speculate that such a delay is a consequence of the specialization of classical RNAi and nuclear RNAi: the former eliminates the immediate threat by degrading the target RNA, and the latter provides long term silencing at the chromatin level, which is critical for retroviruses or retrotransposons. One possible benefit of the delay of nuclear RNAi is to allow the biogenesis of secondary siRNAs (Pak and Fire, 2007; Sijen et al., 2007), which requires mRNA as template for the RdRP activity. If both nuclear RNAi and classical RNAi are fully active at the initial encounter of dsRNA, germ cells may not be able to produce sufficient siRNAs to facilitate silencing inheritance.

**CRISPR provides a unique advantage for studying transgenerational epigenetics.** In this study, we used CRISPR-mediated gene editing to repair a mutation in the *hrde-1* gene, which allowed us to capture the graduate changes in the endogenous targets after the nuclear RNAi machinery is turned on in the subsequent generations. This approach is analogous to heat or small molecule-inducible gene expression, but without the need to change the native promoter or protein sequence of the inducible gene. This approach also avoids crossing two strains carrying different genetic and epigenetic backgrounds, which makes this approach highly tractable. (For example, there is no need to distinguish different epi-alleles.)

Our study showed that, after repairing *hrde-1*, wild type animals fully restored silencing at the endogenous targets in the third generation. Our current implementation of CRISPR does not allow us to examine the first and second generations after the HRDE-1 activity is restored. The one-generation delay observed for the exo-dsRNA-triggered nuclear RNAi raises a possibility that the establishment of silencing at the endogenous targets may also have a transgenerational delay even in a wild type H3K9 HMT background.

CRISPR-mediated genome editing provides a powerful tool for biomedical research and curing genetic diseases. Our study provides a cautionary example that repairing a mutated gene may not immediately restore the normal function expected for the wild type allele. Therefore, epigenetic effects must to be considered when editing genes, especially when chromatin factors and epigenetic pathways are involved.

## Methods and Material

### Strains

Bristol strain N2 was used as a standard wild type strain. This study used the following mutations: LGI: *set-32(ok1457), set-32(red11)*; LGIII: *met-2(n4256), set-25(n5021), hrde-1(tm1200).* The *set-32(red11)* alleles contains a non-sense mutation, followed by a frame shift in the third exon, and was generated in this study using the CRISPR-mediated genome editing procedure described in (Arribere et al., 2014; Paix et al., 2015). Genotyping primers and other relevant sequences are listed in Table S1. Non-RNAi worms were cultured on NGM plates with OP50 *E.coli* as the food source (Brenner, 1974). Worms were cultured at 20°C for all of the experiments conducted in this study. Synchronized young adult animals were ground in liquid nitrogen by mortar and pestle and stored at −80°C.

### *oma-1* RNAi experiments

*oma-1* RNAi experiments were performed as described previously (Kalinava et al., 2017; Timmons et al., 2001) with the following modifications. Two schemes of *oma-1* RNAi were used in this study. To examine the onset of RNAi, synchronized L1 larvae were released onto *oma-1* RNAi plates which contained *oma-1*-dsRNA-expressing E. coli. The young adult animals are referred to as F1(dsRNA+). To obtain worms of F2(dsRNA+) or with extended generations of dsRNA exposure, eggs from dsRNA+ adult animals were collected and hatched in M9 buffer. L1s were released onto *oma-1* RNAi plates for another generation of dsRNA exposure. For the *oma-1* heritable RNAi and H3K9me3 ChIP experiments in Fig. S2, worms of mixed developmental stages were first cultured on *oma-1* RNAi plates continuously for 9-10 days starting with approximately 5 L3-L4 worms, followed by one round of synchronized culture on *oma-1* RNAi plates to collect young adults.

### Gene conversion to repair *hrde-1* mutation using CRISPR-Cas9

We repaired the *hrde-1(tm1200)* mutation to the wild-type sequence using a *hrde-1* targeting Cas-9 ribonuclease complex and PCR fragment as the repair template (relevant sequences are listed in Table S1). A dpy-10(*cn64*) co-conversion marker was used (Arribere et al., 2014; Paix et al., 2015). 10-15 adult hermaphrodite animals (P0s) of each strain were injected. We screened 48-96 F1s animals using single-worm PCR from broods of injected animals with high frequencies of roller F1s. This yielded with 2-8 F1s bearing potential gene conversion events (heterozygotes) identified by the size of PCR products. For each putative F1 hit, 12-24 F2 worms a wild type copy of *dpy-10* were individually transferred to plates. After laying eggs, the F2 worms with homozygous wild type *hrde-1* sequences were identified by PCR and Sanger sequencing. We refer to the CRISPR/Cas-9-generated wild type *hrde-1* sequence as *hrde-1*(+^R^) to distinguish it from the naive wild type *hrde-1* sequence, referred to as *hrde-1(+)*. F3, F4, F5, F10, and F20 post-conversion adults were collected either by hand picking (F3 and F4) or synchronized culture (F5 and later generations). The control *hrde-1(+)* samples (WT, *set-32, met-2 set-25, set-32; met-2 set-25)* and *hrde-1(−)* samples *(*hrde-1, set-32; hrde-1, met-2 set-25 hrde-1* and *set-32; met-2 set-25 hrde-1*)* were collected by picking adult animals, identically to the F3 and F4 samples of the *hrde-1(+^R^)* samples.

### Total RNA extraction

For samples of pooled whole worms, TRIzol reagent (Life Technologies) was added to the frozen sample of approximately 20 worms in the M9 buffer. To ensure break down of worm bodies, we used 3-4 cycles of freeze-thawing in TRIzol, then performed total RNA extraction according to the manufacture’s protocol. This procedure yielded 1-3 μg of total RNA for each sample.

### mRNA and pre-mRNA qRT-PCR

Total RNA extraction was performed using the TRIzol Reagent. 1 μg of total RNA was used for the first strand cDNA synthesis with SuperScript III RT kit (Life Technologies) and oligo-dT as the primer for mRNA qRT-PCR and the random hexamers as the primer mix for pre-mRNA qRT-PCR.

qPCR was performed using KAPA SYBR FAST Universal 2× PCR Master Mix (KAPA Biosystems) on a Mastercycler EP Realplex realtime PCR system (Eppendorf) according to the manufacturer’s instructions. qPCR primers are listed in Table S1. Each sample was processed in triplicate. Reported values for the fold change of mRNA and pre-mRNA were calculated using ΔΔCT analysis. *tba-1* was used as a reference gene.

### High-throughput sequencing

Chromatin immunoprecipitation (ChIP): Pol II and H3K9me3 ChIP-seq was performed as described previously (Ni et al., 2014). The anti-RNA Pol II S2 antibody (ab5095, Abcam) and anti-H3K9me3 antibody (ab8898, Abcam) were used.

RNA-seq: Ribosomal RNA (rRNA) was removed from the total RNA using RNase H and anti-rRNA oligo mixture (total of 110, listed in Table S2). The rRNA-removal procedure was adopted from (Frokjaer-Jensen et al., 2016). Briefly 1.25 μg of anti-rRNA oligos were mixed with 0.5 μg total RNA in 1x Hybridization Buffer (100 mM Tris-HCl pH 7.4, 200mM NaCl) in a final volume of 8 μl. The sample was denatured at 95°C for 2 minutes, and then cooled at −0.1 °C/sec to 45°C. Equal volumes of Hybridase Thermostable RNase H (5U/μl,Epicentre) and 10x RNaseH Reaction Buffer (500 mM Tris-HCl pH 7.4, 1 M NaCl, 200 mM MgCl_2_) were mixed and preheated to 45°C before use. 2 μl of the enzyme mix was added to each reaction mix, which was then incubated at 45°C for 1 hr. To remove DNA oligos, 5 μl TURBO DNase buffer, 3 μl TURBO DNase (Invitrogen), and 32 μl diH2O were added to the reaction mix, followed by 37°C incubation for 30 minutes. To purify RNA, 300 μl STOP solution (1 M ammonium acetate, 10 mM EDTA) was added to the reaction mix, followed by phenol/chloroform (1:1) extraction and ethanol precipitation. The resulting RNA (without poly(A) selection) was used to prepare the RNA-seq library as described previously (Ni et al., 2014).

siRNA-seq: small RNA isolation and RNA-seq library construction were performed as described previously (Ni et al., 2014).

All libraries were sequenced using Illumina HiSeq 2500 platform, with a 50-nt single-end run and dedicated index sequencing. All libraries used in this study are listed in the Table S3.

### Data analysis

Whole genome alignment of the sequencing reads to the *C. elegans* genome (WS190 version) was done using Bowtie (0.12.7) (Langmead et al., 2009). Only perfectly aligned reads were used for data analysis. Reads that aligned to N different loci were scaled by a factor of 1/N. Normalization based on total reads aligned to the whole genome for each library was used for all data analysis, except for the ΔPol II box plot analysis in Fig. S5a, where total reads count of the top 5th percentile of the Pol II signal in corresponding library was used as the normalization factor. The three-region Venn diagram was generated using a web-based software (<u>http://www.benfrederickson.com/venn-diagrams-with-d3.js/</u>). Custom R and python scripts were used in this study. Welch Two Sample t-test was used to calculate all p-values.

## Acknowledgements

We thank Elaine Gavin, Shobhna Patel, Lamia Wahba, and Andy Fire for help and suggestions. Research reported in this publication was supported by the Busch Biomedical Grant and the National Institute of General Medical Sciences of the National Institutes of Health under award number R01GM111752. Some strains were provided by the CGC, which is funded by NIH Office of Research Infrastructure Programs (P40 OD010440). The content is solely the responsibility of the authors and does not necessarily represent the official views of the National Institutes of Health.

---

**Figure S1.**
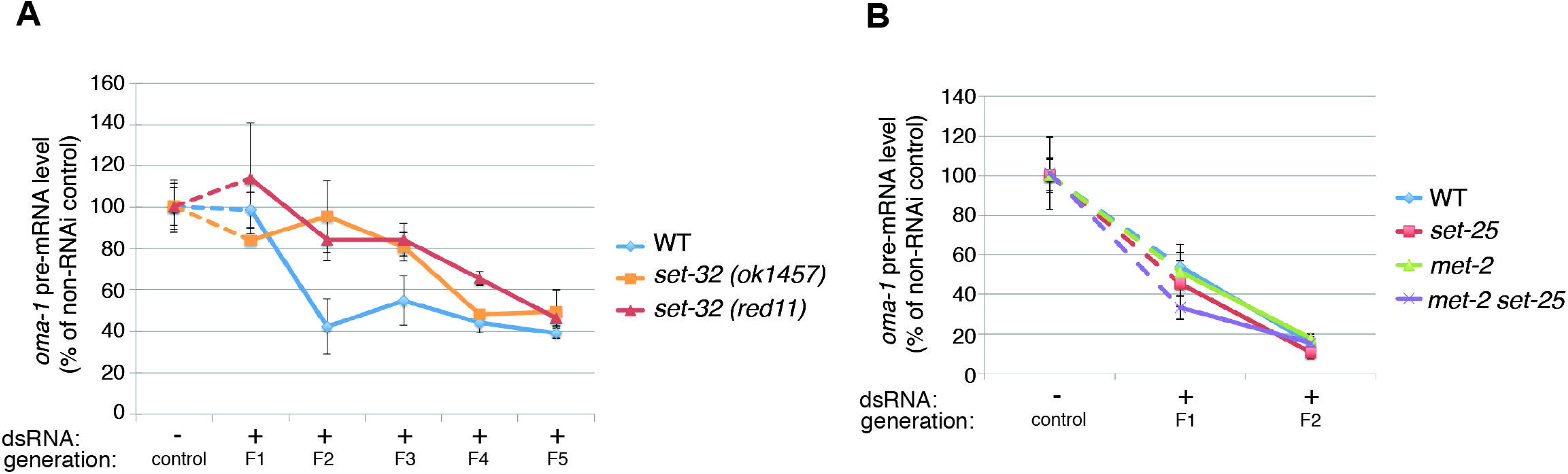
Multigenerational analysis of dsRNA-triggered RNAi at *oma-1.* (A) qRT-PCR analysis of *oma-1* pre-mRNA of the control samples and F1-F5(dsRNA+) samples for wild type and two different *set-32* mutant strains. This is the second biological replica of the experiment shown in Fig. 1C. (B) qRT-PCR analysis of *oma-1* pre-mRNA of the control samples and F1-F2(dsRNA+) samples for WT, *met-2, set-25,* and *met-2 set-25* mutant strains.

**Figure S2.**
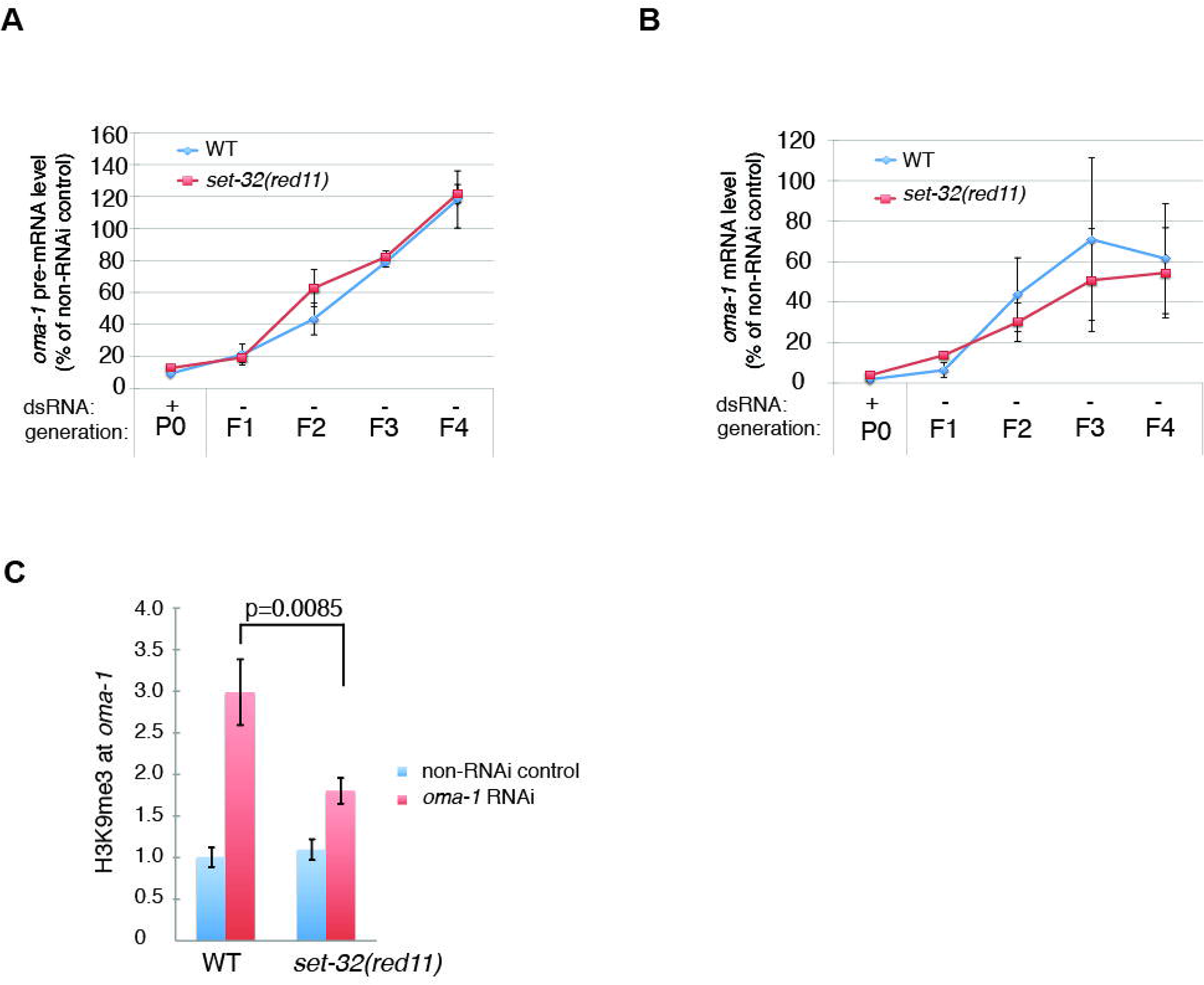
Characterization of the *set-32(red11)* mutant strain. (A and B) Heritable RNAi assay using *oma-1* as the target gene. P0: a population cultured on *oma-1* dsRNA-expressing *E. coli* for approximately 4 generations. F1-F4 progeny of P0 were cultured in the absence of *oma-1* dsRNA. Synchronized adults were collected for all samples. *oma-1* pre-mRNA (A) and mRNA (B) levels were examined by qRT-PCR. Values are expressed as percentage of RNA expression in the non-RNAi control sample of the same genotype. (C) H3K9me3 ChIP-qPCR analysis to examine the exo-dsRNA-induced H3K9me3 at *oma-1* in WT and *set-32(red11)* mutant. OP50-fed adults (control) and adult animals from a population with 4-5 generation *oma-1* dsRNA feeding were used in this assay.

**Figure S3.**
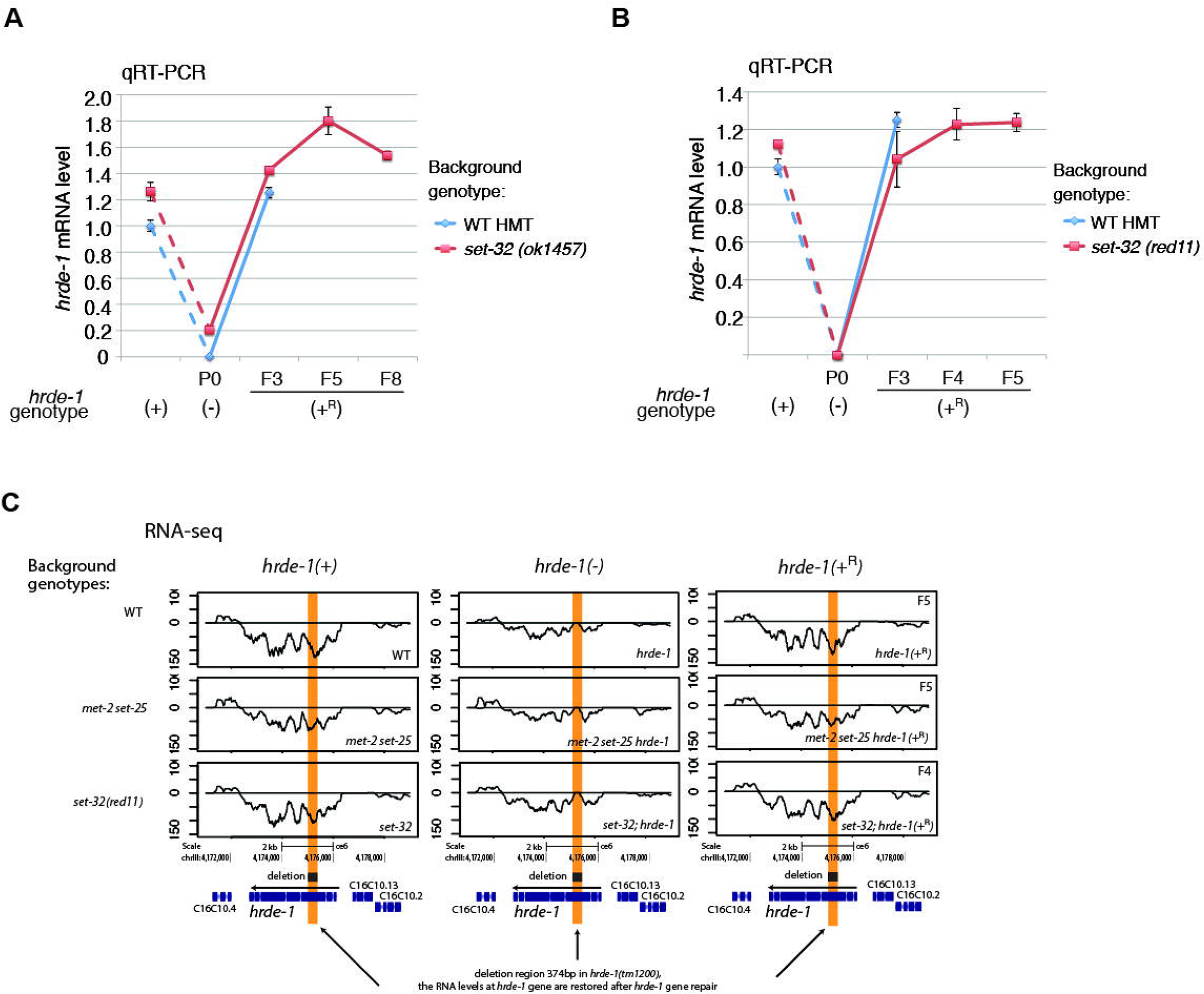
Examining the *hrde-1* mRNA expression in the silencing establishment assay. (A and B) qRT-PCR analysis of *hrde-1* mRNA using primers that are located within the deleted sequence in the *hrde-1(tm1200)* mutation. C. RNA-seq profiles at the *hrde-1* locus. The *hrde-1(tm1200)* deletion is marked by an orange stripe.

**Figure S4.**
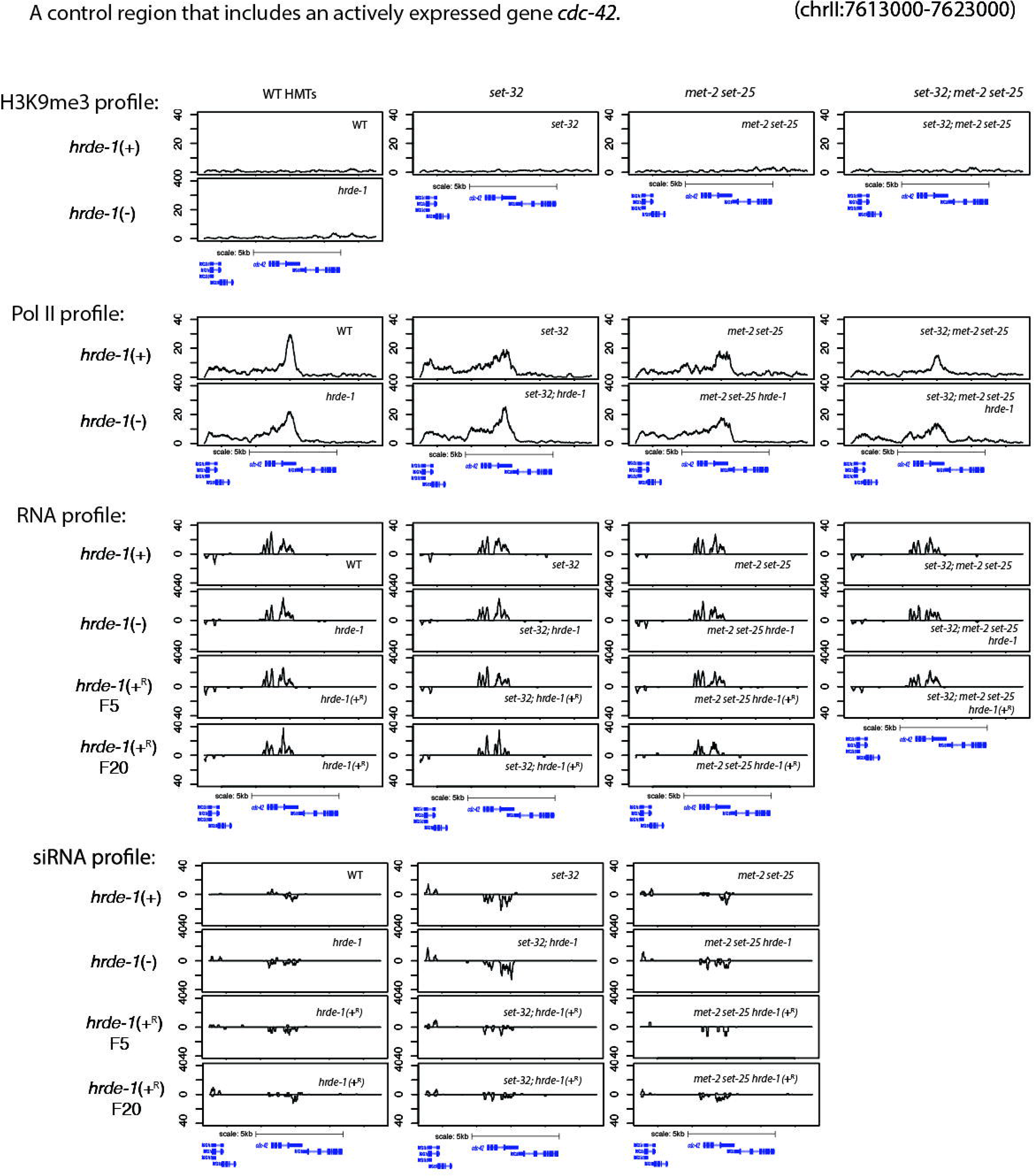
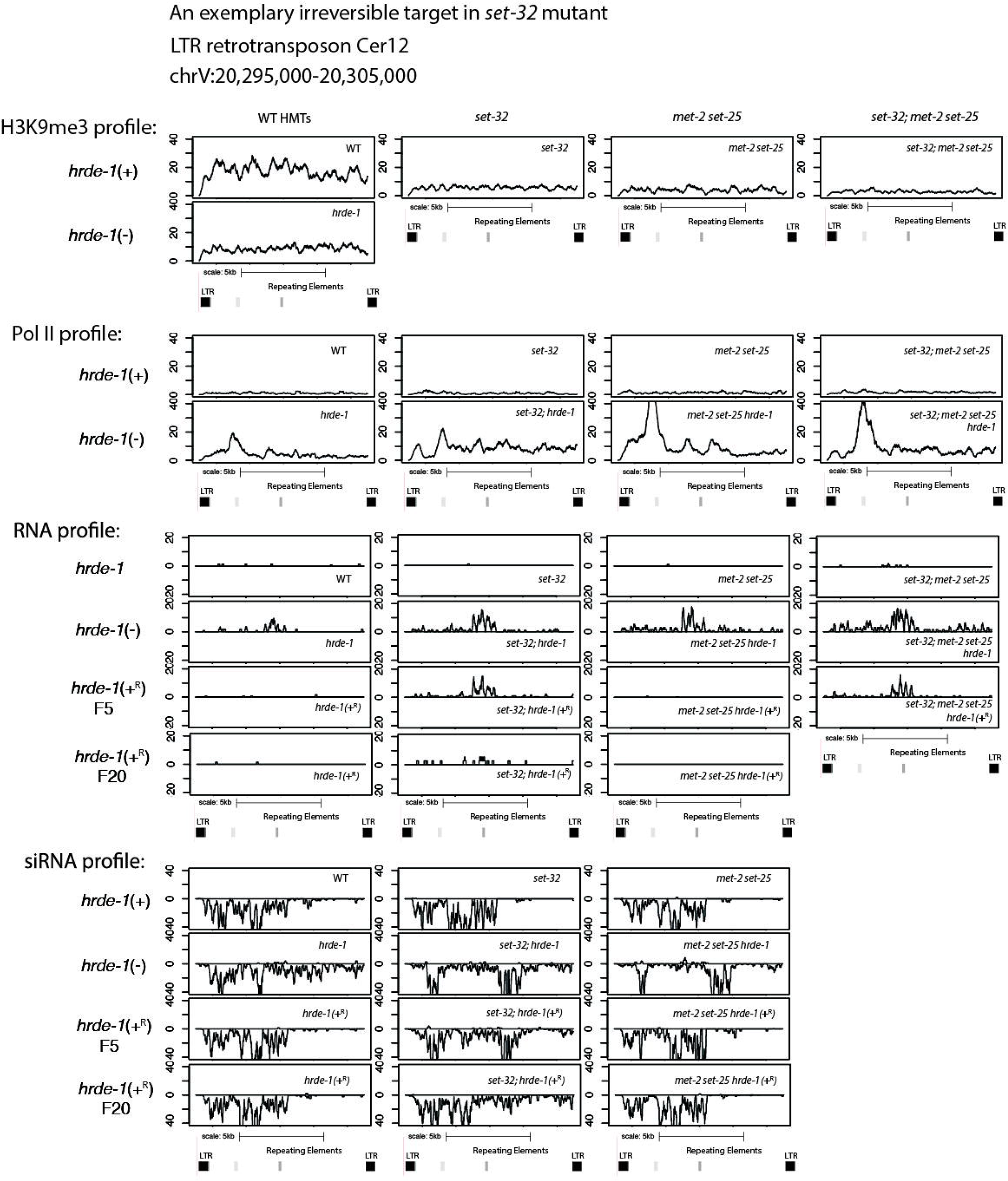
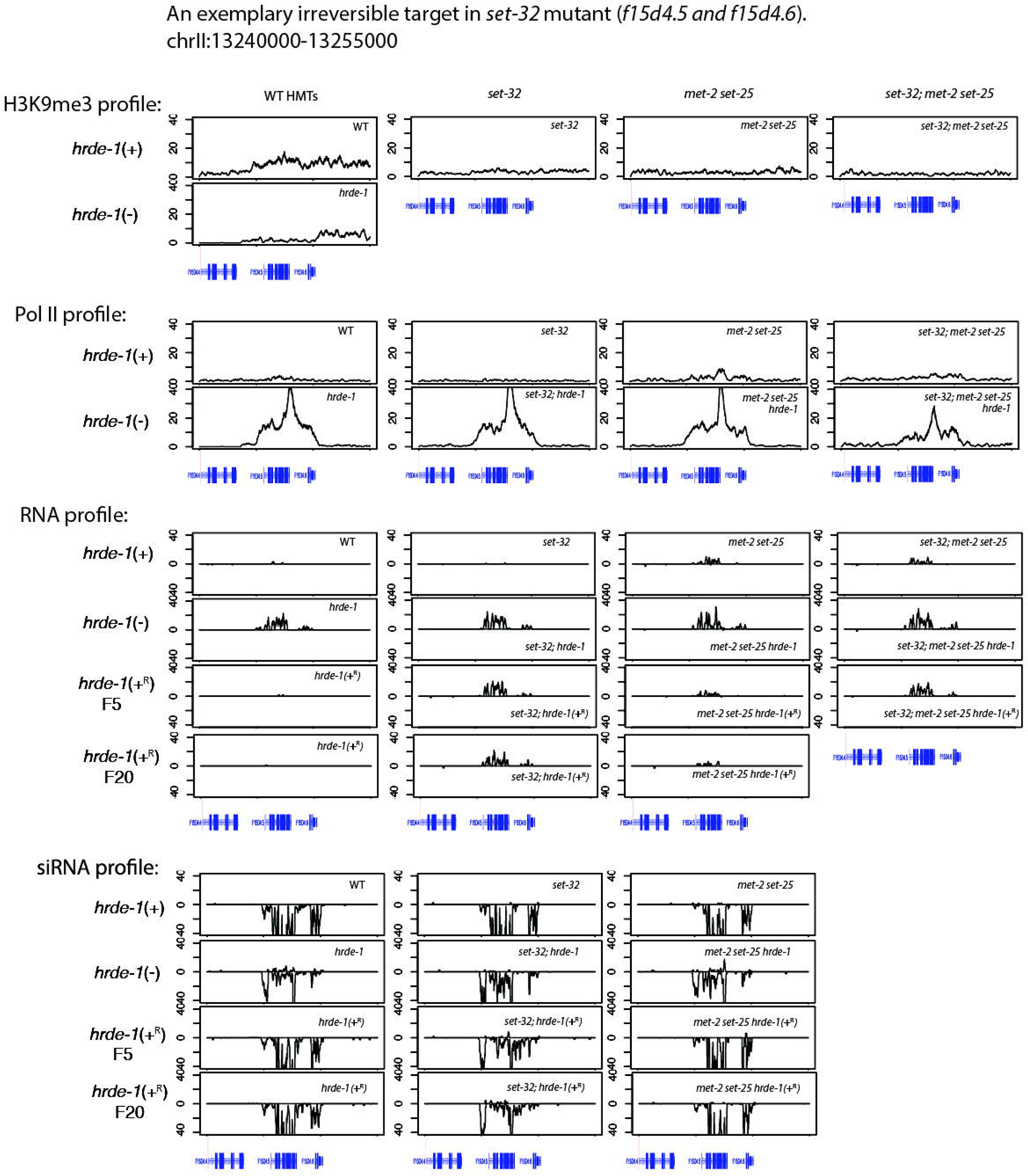
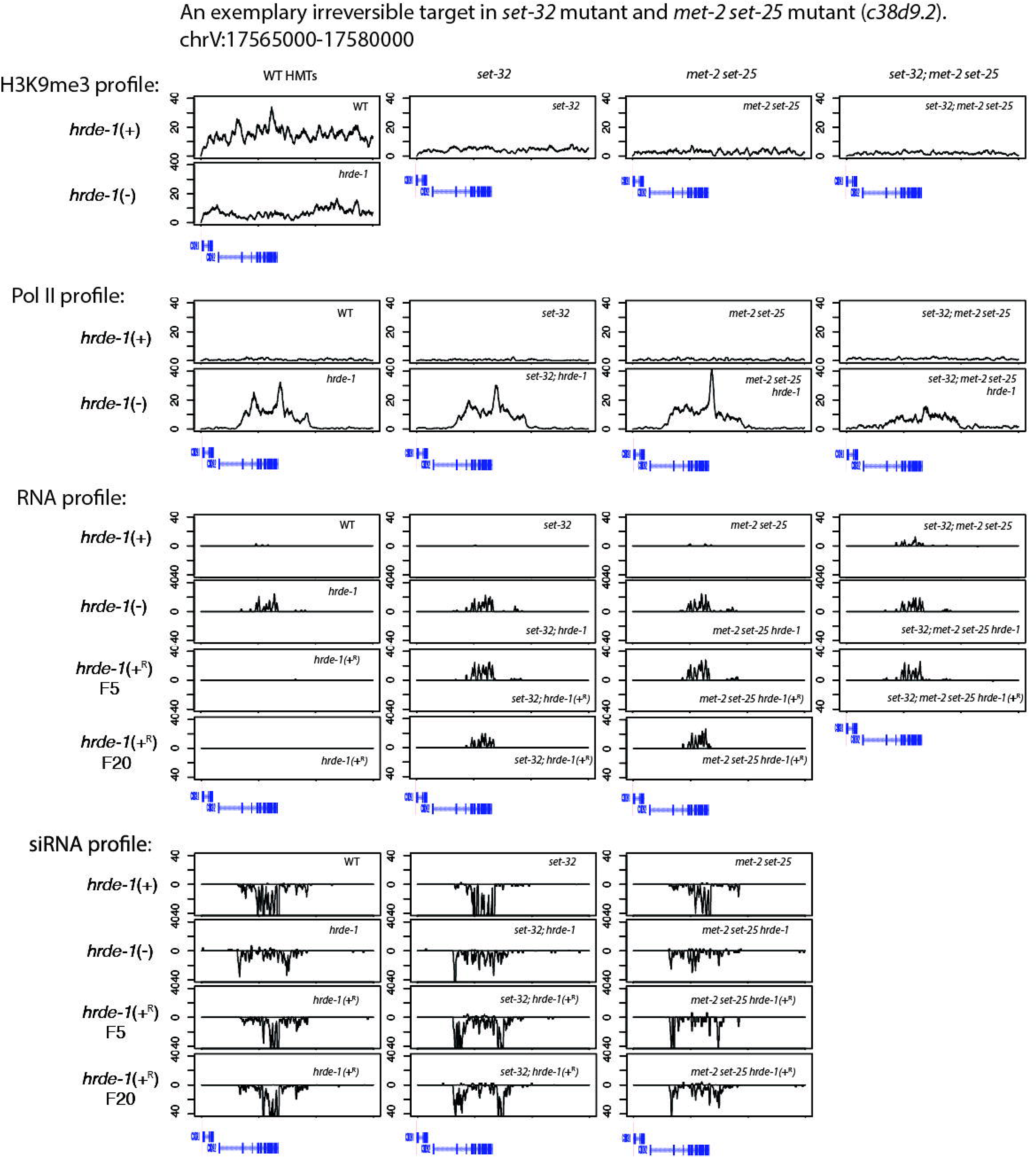
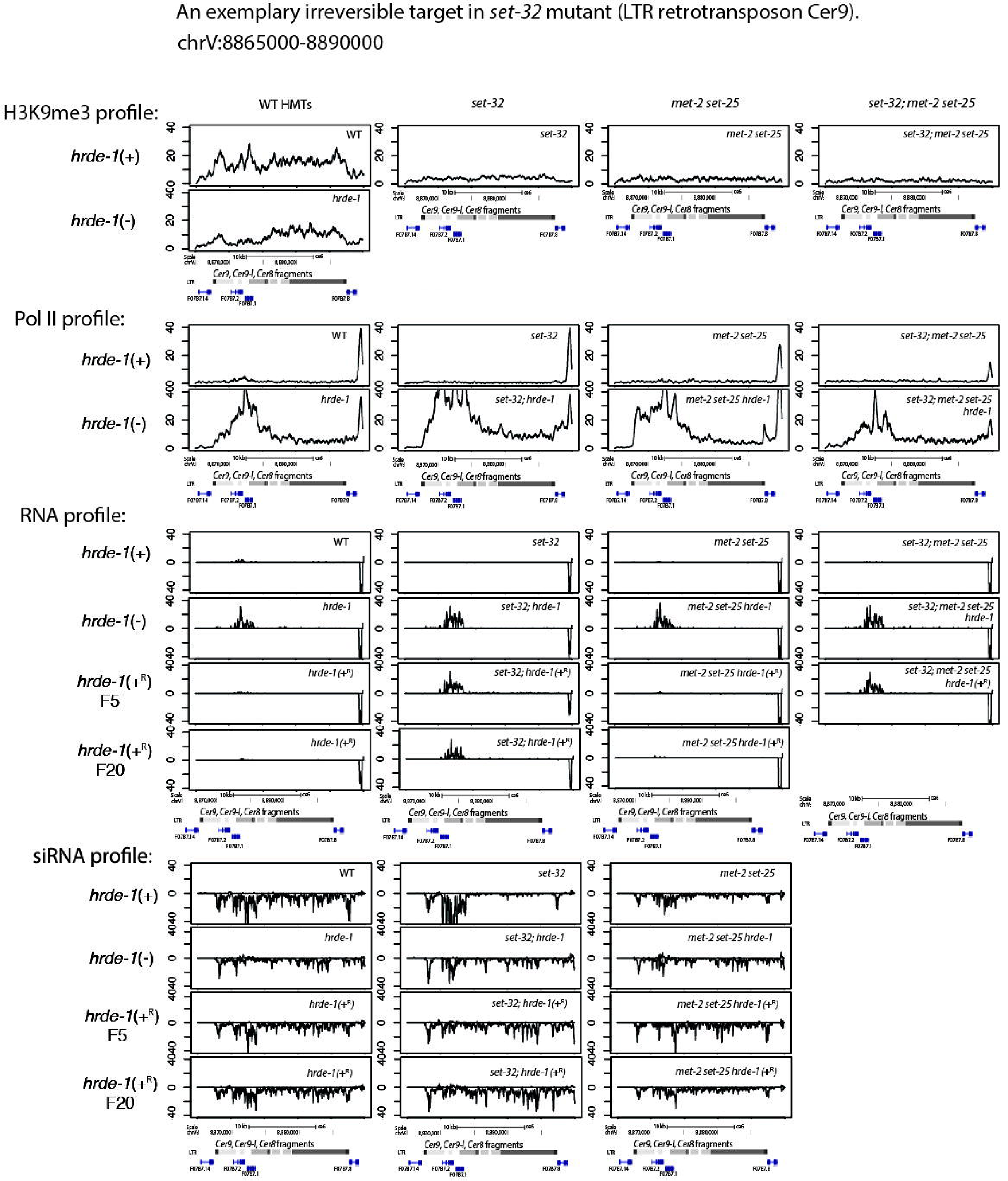
ChIP-seq (H3K9me3 and RNA Pol II [S2 phosphorylated]) and RNA-seq (total RNA and sRNA) profiles at *cdc-42* (a control region, no nuclear RNAi activity) (A) and four exemplary irreversible targets with delayed silencing establishment in *set-32;hrde-1(+^R^)* (B-E). Some of the coverage plots for Cer9 shown in panel E were also presented in Figure 3. The H3K9me3 ChIP-seq data were from our previous study (Kalinava et al., 2017) and the rest of the data were generated in this study. For total RNA and sRNA-seq, the coverages of the “+” and “-”-strand reads are separately plotted (“+”: above the 0-horizontal line, “-”: below).

**Figure S5.**
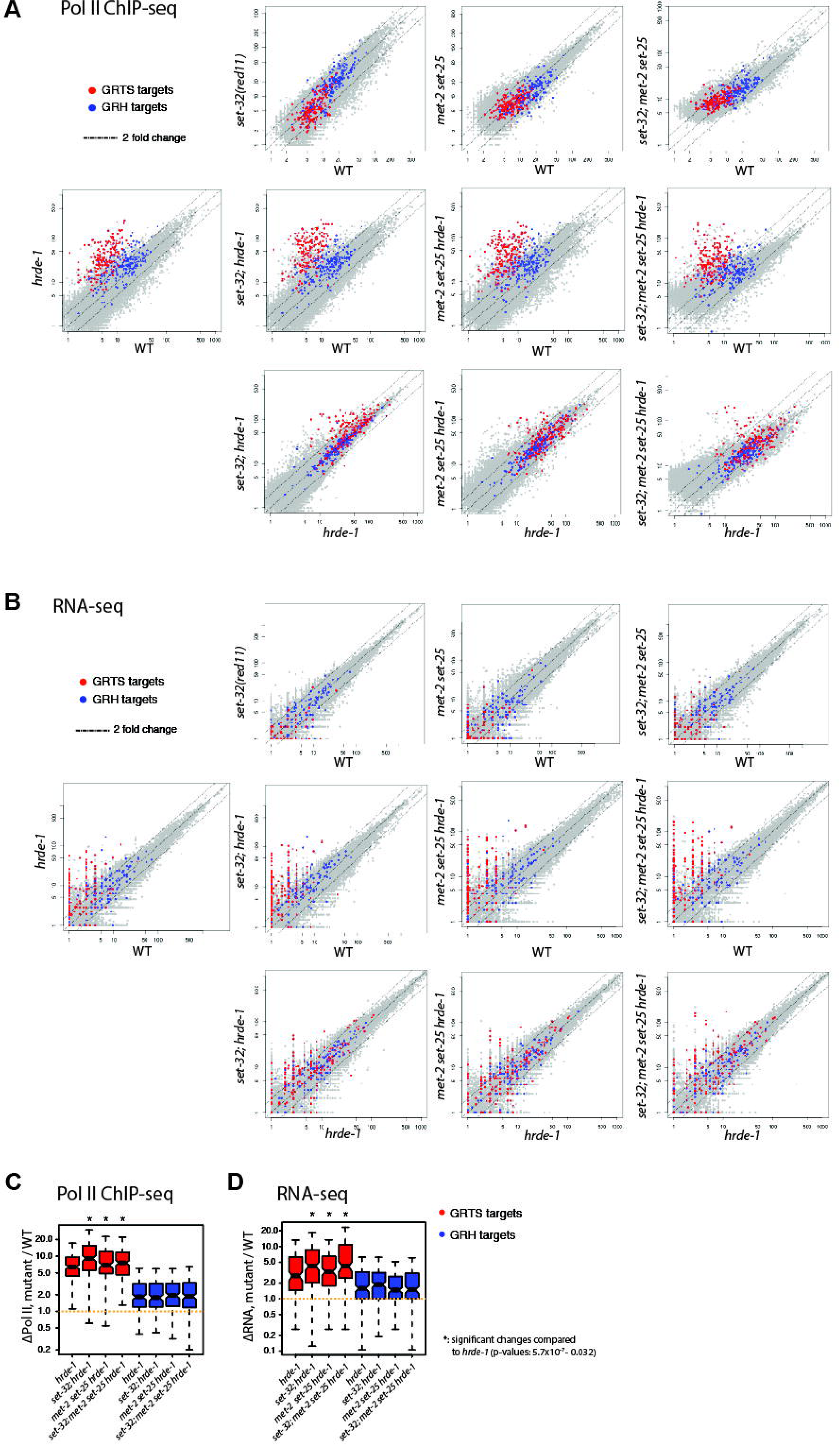
Whole-genome analysis of the enhancement of silencing defects caused by the combination of *H3K9 hmt* and *hrde-1* mutations. (A and B) Boxplot analysis of the changes in Pol II (S2 phosphorylated) occupancy (A) and RNA-expression (B) of GRTS or GRH regions between various mutants or wild type animals.

